# VHUT-cryo-FIB, a method to fabricate frozen-hydrated lamella of tissue specimen for *in situ* cryo-electron tomography

**DOI:** 10.1101/727149

**Authors:** Jianguo Zhang, Danyang Zhang, Lei Sun, Gang Ji, Xiaojun Huang, Tongxin Niu, Fei Sun

## Abstract

Cryo-electron tomography (cryo-ET) provides a promising technique to study high resolution structures of macromolecules *in situ*, opening a new era of structural biology. One major bottleneck of this technique is to prepare suitable cryo-lamellas of cell/tissue samples. The emergence of cryo-focused ion beam (cryo-FIB) milling technique provides a good solution of this bottleneck. However, there are still large limitations of using cryo-FIB to prepare cryo-lamella of tissue specimen because the thickness of tissue increases the difficulty of specimen freezing and cryo-FIB milling. Here we report a new workflow, VHUT-cryo-FIB (Vibratome - High pressure freezing - Ultramicrotome Trimming – cryo-FIB), aiming for efficient preparation of frozen hydrated tissue lamella for subsequent cryo-ET data collection. This workflow includes tissue slicing using vibratome, high pressure freezing, ultramicrotome cryo-trimming, cryo-FIB milling and the subsequent cryo-electron microscopy (cryo-EM). The modification of equipment in this workflow is highly eliminated. We developed two strategies with a special cryo-holder tip or carrier for loading cryo-lamella into side entry cryo-holder or Autoloader catridge. We tested this workflow using the tissue sample of rat skeleton muscle and spinach leaf and collected high quality cryo-ET tilt series, which enabled us to obtain an *in situ* structure of spinach ribosome by sub-tomogram averaging.

## INTRODUCTION

Cryo focused ion beam (cryo-FIB) is now widely used to prepare thin frozen hydrated lamellas of cells for *in situ* structural biology study. The lamella can be milled to 200 ∼ 300 nm thickness suitable for cryo-electron tomography (cryo-ET) without common artifacts caused by cryo-ultramicrotomy ^1,2^. In 2006, Marko et al. first reported vitreous water could be thinned by cryo-FIB and no re-crystallized ice appears in the final cryo-lamella ^3^. A year later, this method was first applied on *E.coli* for the definitive proof of concept ^4^. After that, cryo-FIB method was further applied to eukaryotic cells to study structure of nuclear pore complexe (NPC) in *Dictyostelium discoideum* cell ^1^, architecture of membranous organelle and related protein complexes in *Chlamydomonas* cell^5-7^, thylakoids in cyanobacterium *Synechocystis* ^8^ and the protein sociology around nuclear membrane in Hela cell ^9^. In addition, cryo-FIB was also showed useful in sample preparation for electron diffraction ^10-12^. However, the thickness of sample is limited to cells and crystals that can be well frozen by plunge freezing.

Instead of cells grown in suspension culture or directly on EM grids, the tissue specimen taken from a living body keep at a more close-to-living state and can provide more native structural information. High pressure freezing (HPF) is the unique method currently used for frozen fixation of tissue specimen, and has been widely applied to study ultrastructure of cells and tissues. However, the current procedure of HPF could not produce the frozen specimen with suitable thickness for the subsequent cryo-FIB milling, which restricts the preparation of cryo-lamella of frozen hydrated tissue specimen for *in situ* structural study.

There were few attempts to solve this problem. In 2014, Marko’s group reported a strategy ^13^ that frozen the sample in a carrier using HPF and pretrimmed the carrier by cutting away a substantial portion of the carrier by cryo-ultramicrotome for further cryo-FIB milling. The whole procedure needs a customized intermediate specimen holder (ISH), and a specimen block for cryo-FIB experiment. In addition, the cryo holder in cryo transmission electron microscopy (cryo-TEM) experiment needs to be modified to accommodate the ISH. A year later, Baumeister’s group frozen *C. elegans* worms on a grid using HPF and applied a previously developed cryo-planning method on the surface of the frozen specimen before cryo-FIB milling.^14,15^ However, plenty of customized equipment are still needed. In the same year, Harapin et al.^16^ tried to eliminate the obstacle by changing the traditional cryoprotectant into 2-methylpentane because it is easy to sublime under -150 °C and high vacuum. Thus the sample frozen on the grid is easier exposed, but the following cryo-FIB milling procedure is still difficult to deal with the excessive specimen thickness. All these methods above either need multiple customized equipments or are only suitable for limited kinds of tissue specimen. Therefore, new strategies are demanded to improve the efficiency of using HPF and cryo-FIB for cryo tissue lamella preparation.

Here we report a workflow named VHUT-cryo-FIB (Vibratome - High pressure freezing - Ultramicrotome Trimming – cryo-FIB) to prepare a cryo tissue lamella for high quality cryo-ET data collection. The whole workflow includes fresh tissue sample slicing using vibratome, high pressure freezing, pre-trimming by cryo-ultramicrotome, cryo-FIB milling and final cryo-ET data collection. The whole sample preparation procedure can be performed with limited customized equipments (a modified carrier, a transfer shutter and a modified cryo holder tip) and applied on various kinds of tissue specimen.

## RESULTS

The schematic workflow of VHUT-cryo-FIB is shown in **Fig. 1**. A bulk of tissue immersed in phosphate buffered saline (PBS) is first sliced to 60∼100 μm by vibratome, and then the slice is transferred into a carrier and frozen using HPF. To reduce the overall milling time of cryo-FIB, after HPF a following pre-milling procedure is performed to achieve a preliminary reduction of the specimen thickness from ∼ 100 μm to ∼ 20 μm by cryo-ultramicrotome. Then, cryo-FIB milling is carried on to further reduce the specimen thickness to 200 ∼ 300 nm. The final cryo-lamella can be used for the subsequent cryo-TEM experiment and high quality tilt series data collection.

**Figure 1.**
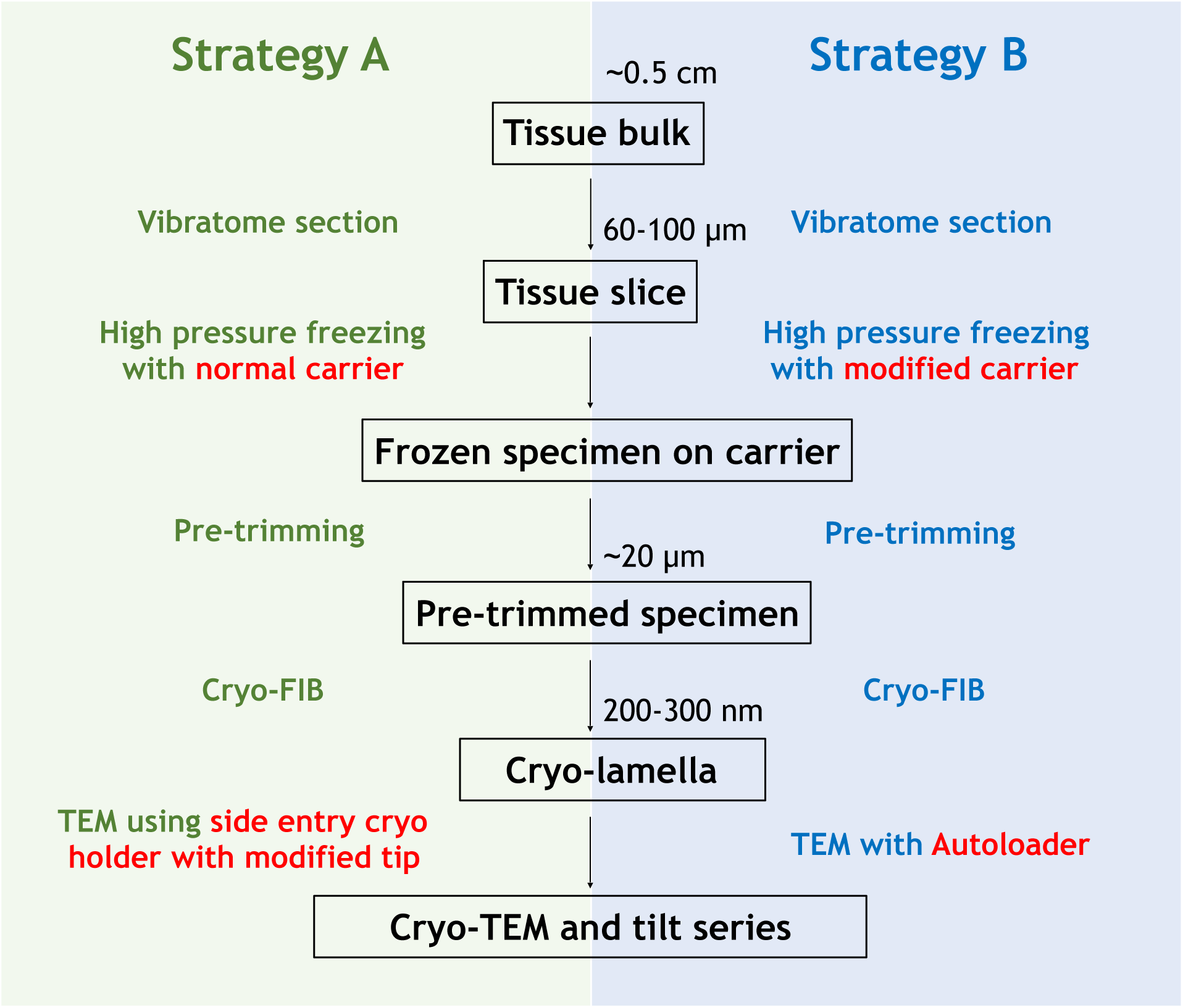
Scheme of VHUT-cryo-FIB workflow. Strategy A is suitable for cryo-TEM using side entry cryo holder and strategy B is suitable for cryo-TEM with FEI Autoloader.

To enlarge the usage of the workflow and limit the requirement of customized equipment, we developed two different strategies to suit different cryo-TEM equipments (**Fig. 1**). Strategy A is suitable for cryo-TEM using side entry cryo holder. In this strategy, a normal HPF carrier (**Figs. 2a and 2b**) and cryo transfer shutter (**Figs. 2c and 2d**) are used, and the tip of a Gatan 910 cryo holder is modified (**Figs. 2e and 2f**). The removable tip has the modified recess in the depth and diameter to accommodate the normal HPF carrier. After loading the pre-trimmed carrier into the tip, a c-clip is inserted into the recess to fix the carrier (**Figs. 2e-2h**). Strategy B is suitable for cryo-TEM equipped with FEI Autoloader (**Supplementary Movie 1**). The only modified devices are the carrier and related accessories used for HPF (**Figs 2i-2n and Supplementary Fig. 1**). We designed an edge of the carrier to match the size of FEI AutoGrid and c-clip, and reduced the depth of the recess from 200 μm (in normal HPF carrier) to 100 μm to improve freezing quality. Assembling with the special designed auxiliary ring, the modified carrier can be frozen and trimmed just as the normal HPF carrier. Then the auxiliary ring is removed and the pre-trimmed carrier is trapped in an FEI AutoGrid (**Figs. 2o and 2p**) for loading into either cryo-FIB chamber (**Fig. 2q**) or cryoTEM column (**Fig. 2r**).

**Figure 2.**
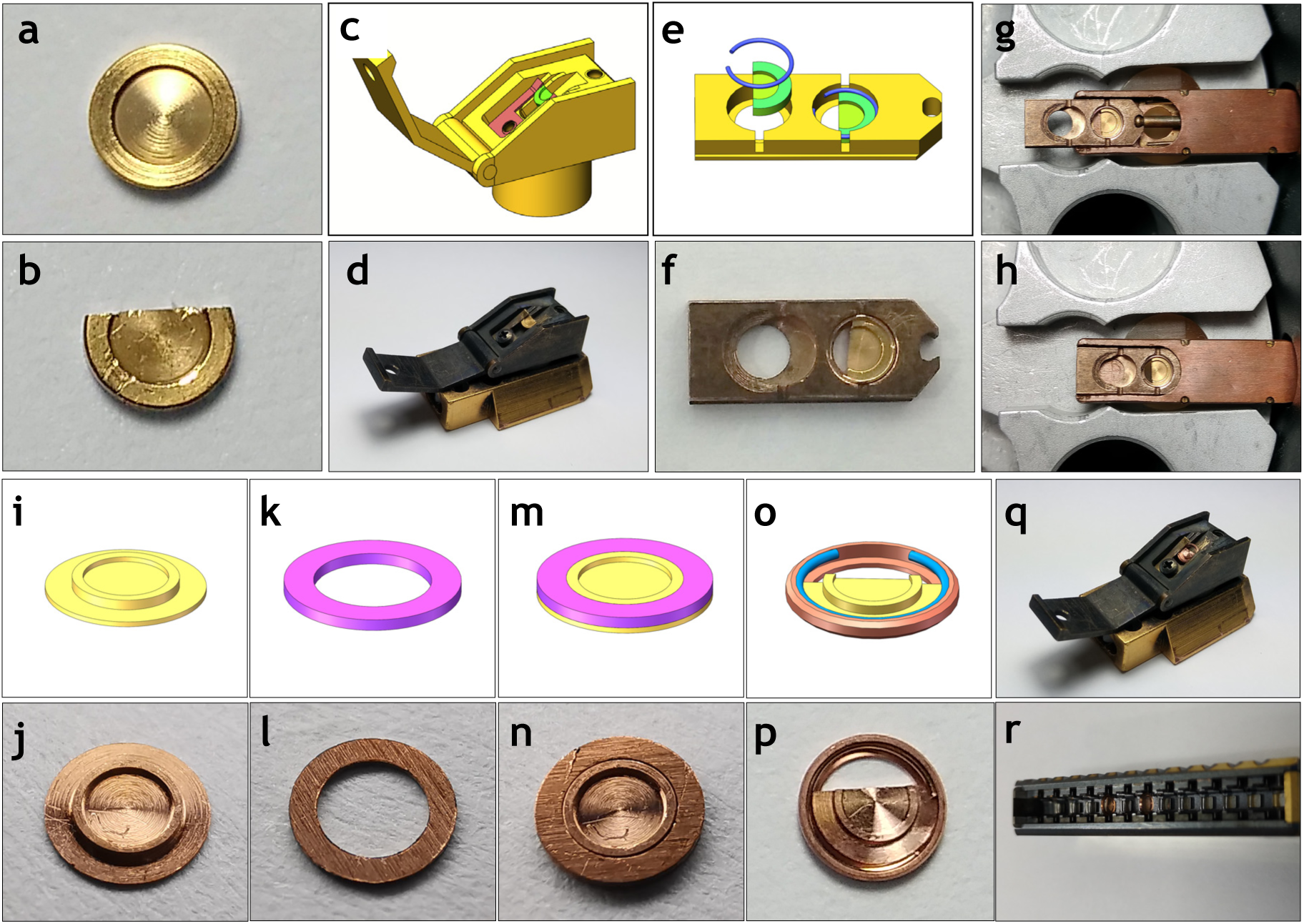
Design and application of customized tools. (a) Carrier used for high pressure freezing in strategy A. (b) The remaining part of the carrier after pre-trimming. (c) Design of the FIB transfer shuttle for loading the pre-trimmed carrier. (d) The real shuttle used in the workflow. (e) Design of the modified tip for Gatan 910 cryo holder and the way of sample loading. (f) The real tip used in the workflow with a sample loaded. (g) Mounting of the modified tip onto the Gatan 910 cryo holder. (h) The cryo holder ready for loading into TEM. (i) Design of the modified carrier used in strategy B. (j) The real modified carrier used in the workflow. (k) and (i) Design and the real picture of the auxiliary ring. (m) and (n) The assembly of the modified carrier and the auxiliary ring and their real picture. (o) and (p) Mounting the trimmed carrier into AutoGrid and the real picture. (q) The transfer shuttle with the trimmed carrier loaded. (r) Loading the AutoGrid with the trimmed carrier into the cassette of FEI Titan Krios.

Vibratome can generate a tissue slice with a thickness of 60 ∼ 100 μm and the slice was further high pressure frozen in a carrier to get vitrified. Then the vitrified specimen together with the carrier were loaded into the chamber of a cryo-ultramicrotome for cryo pre-trimming. One third to half of the carrier together with the ice was trimmed away, leaving a straight edge (**Figs. 2b and 2p**). Moreover, at the straight edge, the carrier and ice on the surface were further trimmed 30 μm deep to leave a 20 μm thick specimen platform exposed outside the edge (**Fig. 3a, 3b and 3c**). Trimmed carrier can be mounted directly into a cryo-FIB transfer shuttle and loaded for cryo-FIB milling (**Figs. 2c, 2d, and 2q**). The platform can be easily found in SE (secondary electron) images produced by both electron beam (SEM mode) and gallium ion beam (FIB mode) (**Figs. 3b and 3c**). Cryo-FIB milling was preformed along the direction of diameter of the carrier and vertical to the straight edge (**Figs. 3c, 3d and 3e**). A slight tilt angle of ion beam is set to avoid the block by FEI AutoGrid. A slight modification we made during milling is to jag a gap at one side of the cryo-lamella (**Fig. 3e**). This could facilitate the release of internal stress in the sample which would cause the bending of cryo-lamella (**Supplementary Fig. 2**). After cryo-FIB milling, a thin cryo-lamella can be observed by SE images in both SEM and FIB modes (**Fig. 3d and 3e**). To be noted that, in our strategies, cryo-FIB milling can be performed at multiple positions on a single platform (**Fig. 3f**), which increases the usage efficiency of one frozen tissue specimen.

**Figure 3.**
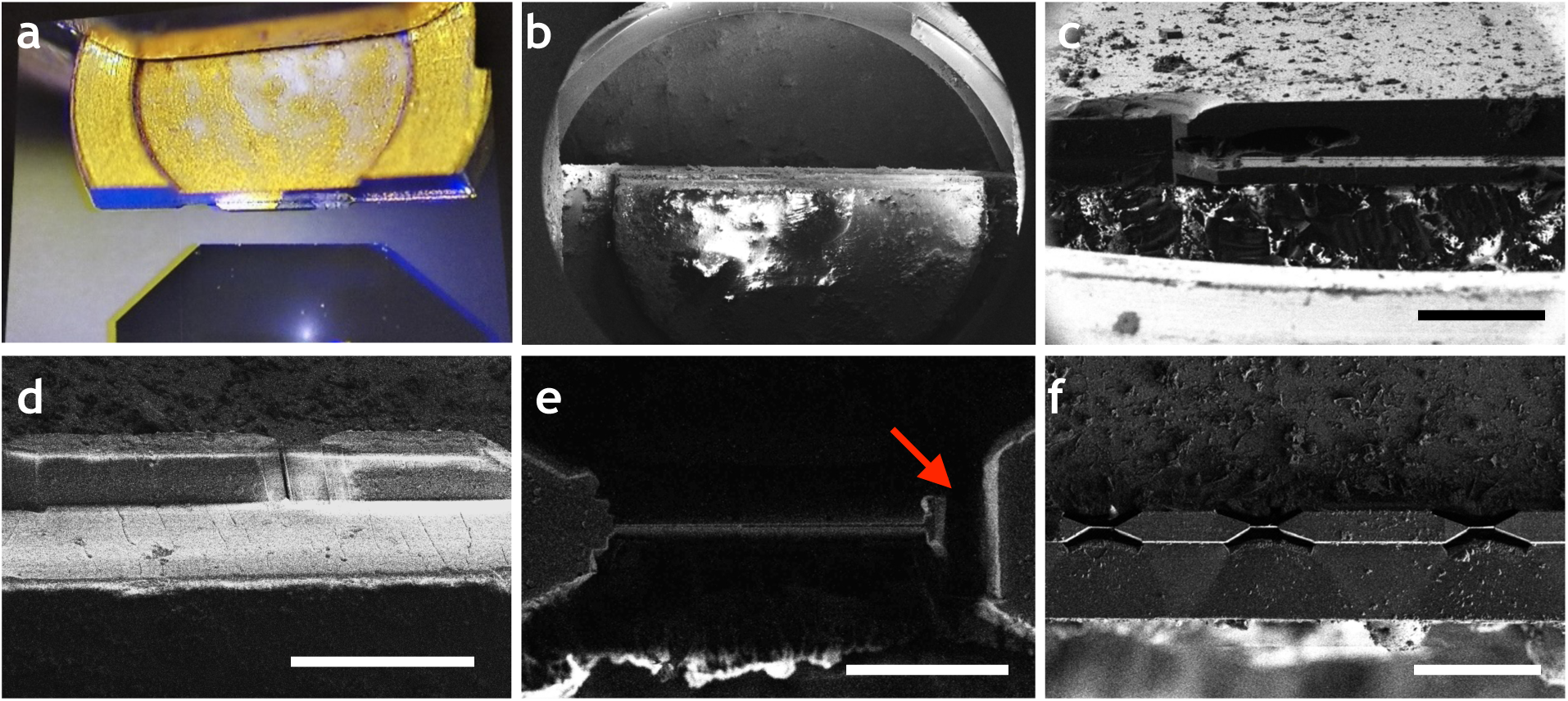
Cryo-FIB milling of the pre-trimmed sample. (a) Picture of a pretrimmed carrier in the cryo-chamber of a cryo-ultramicrotome. (b) SE micrograph of the exposed sample platform in SEM mode. (c) SE micrograph of the exposed sample platform in FIB mode. (d) SE micrograph of the milled cryo-lamella taken in SEM mode. (e) SE micrograph of the milled cryo-lamella in FIB mode. A jag at one end of the cryo-lamella is indicated with the red arrow. (f) Multiple milling positions on a single sample platform. Scale bars, 200 μm in (c), 100 μm in (d), 10 μm in (e), and 100 μm in (f).

Next, the whole carrier is mounted onto a side entry cryo holder or FEI Autoloader for cryo-TEM imaging (**Figs. 2g, 2h and 2r**). For the cryo-lamella imaged by 200kV Talos F200C with Gatan 910 cryo holder, the panorama of cryo-lamella is shown in **Fig. 4a**. The image indicates a well vitrified cryo-lamella with a suitable thickness for tilt series data collection. The electron diffraction pattern (**Fig. 4b**) indicates that the cryo-lamella was kept in a vitreous state. An image taken at higher magnification reveals the details of the specimen (**Fig. 4c**). The specimen we used here is the rat skeleton muscle, and we can clearly see the parallel Z-line and M-line arranging regularly. Beside the myosin filament that is perpendicular to the Z-line, mitochondria and sarcoplasmic reticulum (SR) can also be identified. We then tried to collect tilt series data of the cryo-lamella (**Supplementary Movie 2**). 3D reconstruction showed more structural details of the specimen including density of protein complexes at the intermembrane space of SR (**Fig. 4d and Supplementary Movie 3**). The total area of the cryo-lamella is about 150 μm^2^, which suggests several tilt series can be collected from one cryo-lamella.

**Figure 4.**
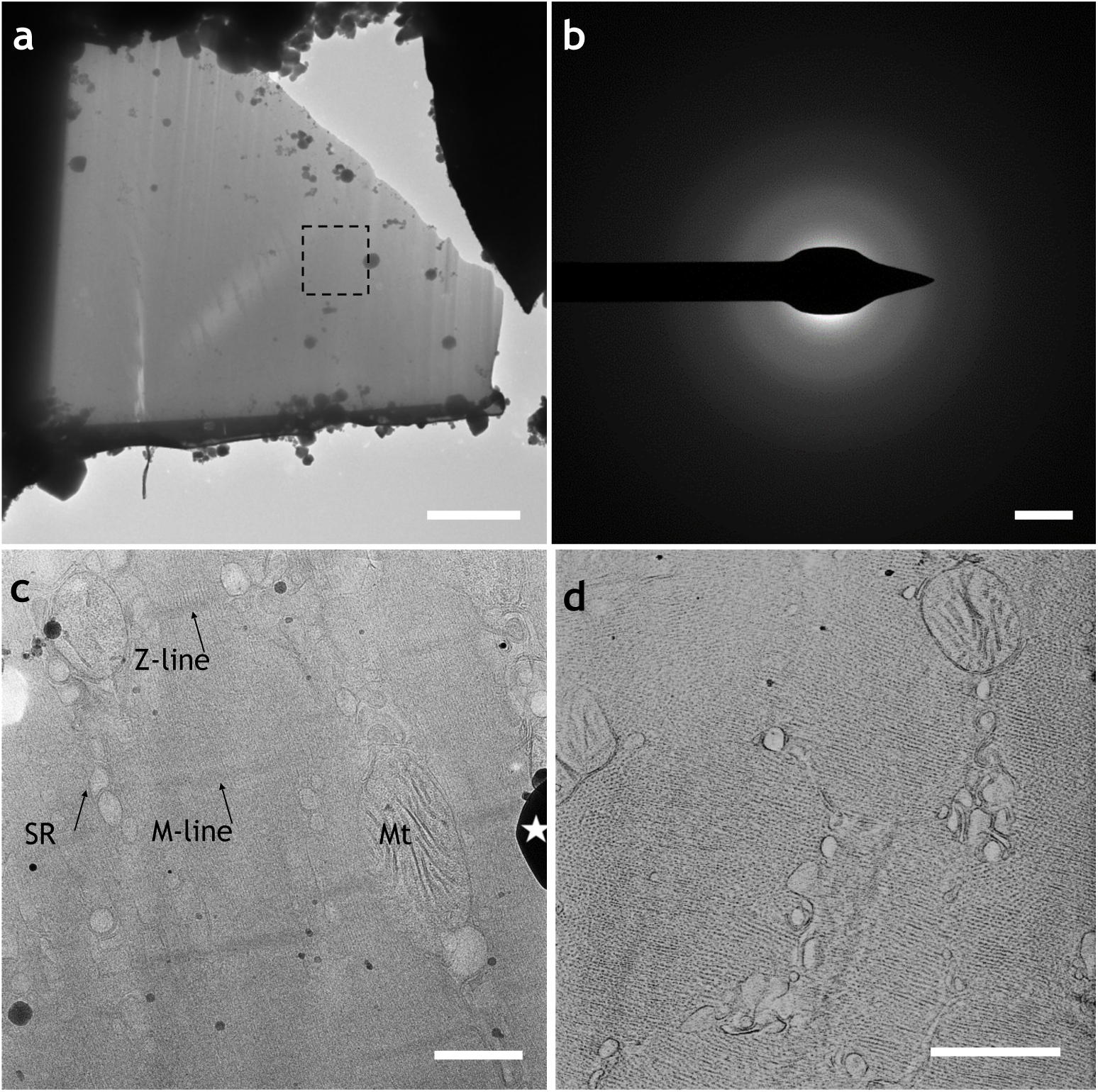
Cryo-lamella of rat skeleton muscle sample made via strategy A. Cryo-electron micrograph of the cryo-lamella in low magnification. (b) Electron diffraction of the cryo-lamella in (a). (c) Cryo-electron micrograph of the cryo-lamella in high magnification at the position in (a). Z-line, M-line, SR, (sarcoplasmic reticulum) and Mt (mitochondria) are labeled. (d) One representative slice of the reconstructed tomogram of the cryo-lamella. The tomogram was reconstructed using ICON. Scale bars, 5 μm in (a), 1.2 1/nm in (b), and 500 nm in (c) and (d).

Cryo-lamella made via strategy B is loaded into a 300kV FEG cryo electron microscope FEI Titan Krios (ThermoFisher Scientific) for further imaging. The low magnificent micrograph also indicates a high-quality cryo-lamella that has a large area suitable for cryo-ET data collection (**Fig. 5a**). For the spinach leaf tissue specimen, the high magnification micrograph shows detailed native ultrastructure of chloroplast with well-ordered thylakoids (**Fig. 5b**). The equidistant membrane structure in granum is easy to identify and the connection between neighbor granums can also be seen clearly. For the rat skeleton muscle cryo-lamella (**Fig. 5c**) made via this strategy, the myosin, SR and mitochondria can also be identified (**Fig. 5d**).

**Figure 5.**
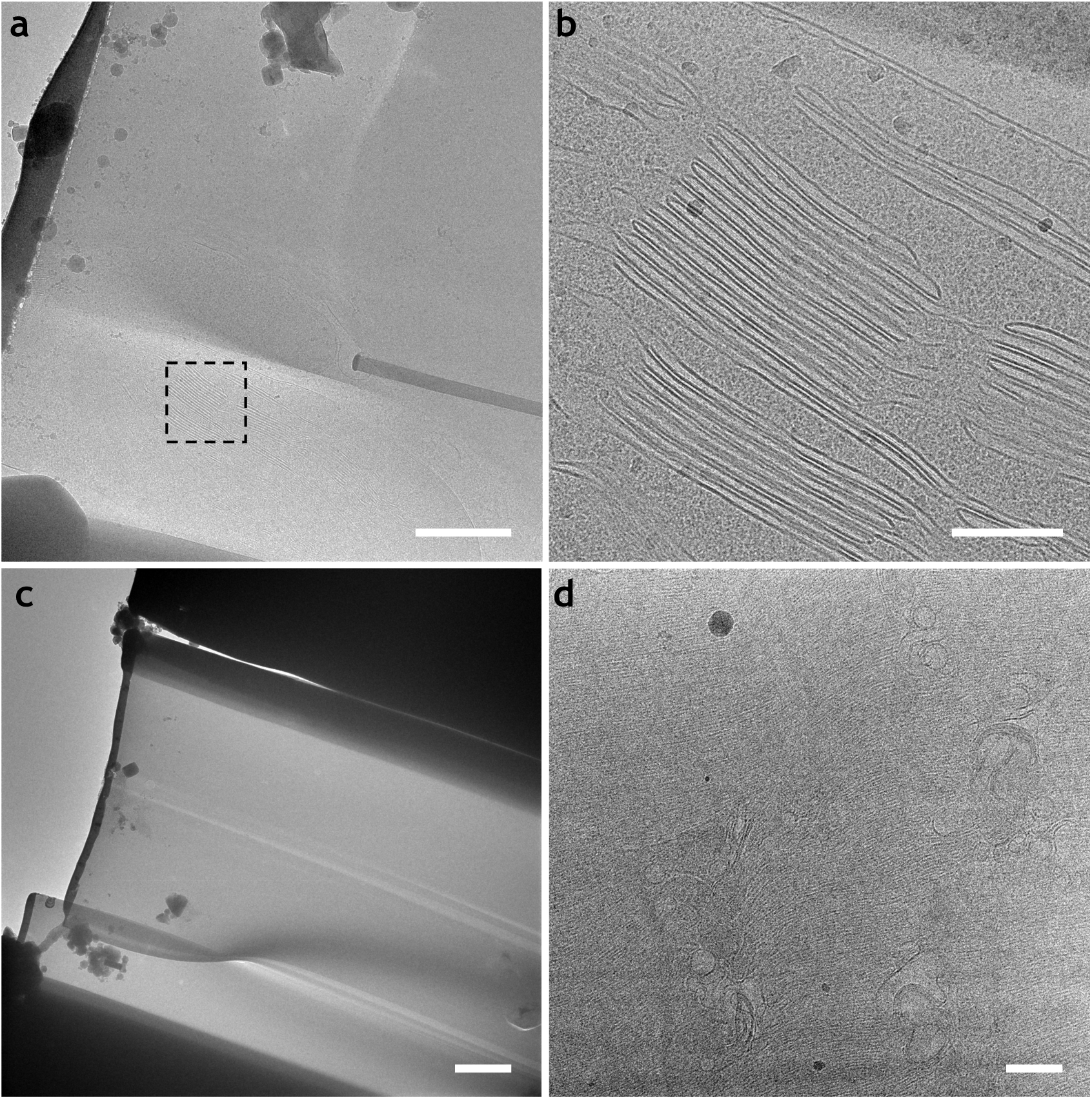
Cryo-lamellas of tissue sample made via strategy B. (a) Cryo-electron micrograph of the cryo-lamella of spinach leaf in low magnification. (b) Cryo-electron micrograph of the cryo-lamella of spinach leaf in high magnification at the position in (a). (c) and (d) Cryo-electron micrographs of the cryo-lamella of rat skeleton muscle in low (c) and high magnification (d). Scale bars, 1 μm in (a), 200 nm in (b), 2 μm (c), and 500 nm in (d).

We then collected tilt series cryo-ET data of the spinach leaf specimen (**Supplementary Movie 4**). In order to avoid beam blocking by the ice behind the cryo-lamella during tilting, the rotation axis was set perpendicular to the path of FIB beam (**Supplementary Fig. 3a**). Considering the incident angle of FIB beam, a single focus position could not report the accurate defocus value during data collection. Therefore, we adopted two focus steps at each side of the recording area and took the averaged defocus value as reference for autofocusing (**Supplementary Fig. 3a**). This approach yielded a stable defocus for each tilt exposure (**Supplementary Fig. 3b**). The reconstructed tomogram of spinach leaf cryo-lamella exhibits good contrast that allows direct visualization and identification of organelle and molecular complexes (**Fig. 6a and Supplementary Movie 5**). Fibrous densities were observed in cell wall, indicating the network formed by cellulose and hemicellulose (**Figs. 6a and b**). Near the cell wall, a chloroplast was identified by its double layer membrane and classic granum (**Figs. 6a and b**). In addition to the membranous structure, the remaining matrix in the chloroplast was filled with granular densities, representing variety kinds of protein complexes. In the cytoplasm between cell wall and chloroplast, abundant ribosomes were found and some of them were adhering with mRNAs to form polyribosomes. We picked out these ribosome particles and performed sub-tomogram averaging. An averaged map of spinach ribosome was obtained with a resolution of of 34 Å according to the gold standard FSC_0.143_ criteria (**Supplementary Figs. 4a and 4b**). The large and small subunits of spinach ribosome can be clearly identified (**Fig. 6c**) and its general structure is similar to the 80S human ribosome (PDB code 6EK0) (**Supplementary Fig. 4c**).

**Figure 6.**
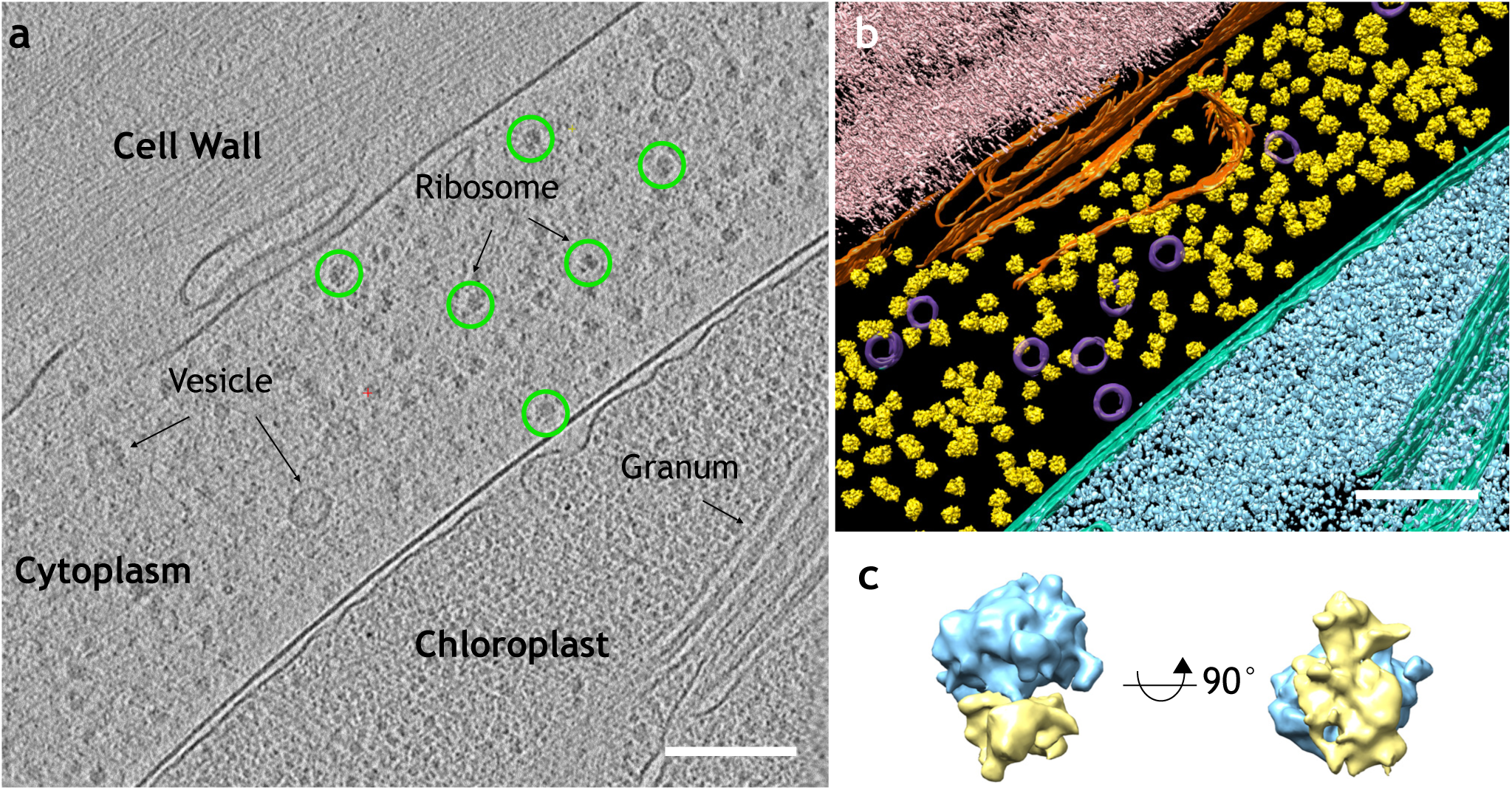
Cryo-ET of spinach leaf cryo-lamella and the sub-tomogram averaging analysis. (a) One representative slice of the reconstructed tomogram of spinach leaf cryo-lamella. The native ultrastructures including cell wall, cytoplasm, chloroplast, vesicles, ribosomes and granum are labeled accordingly. (b) Segmentation of the reconstructed tomogram. Yellow, ribosomes; Light pink, cell wall; Orange, cytoplasmic membrane; Magenta, vesicles; Green, chloroplast membrane; Cyan, chloroplast matrix. (c) *In situ* structure of spinach ribosome from sub-tomogram averaging analysis. Its large subunit is labeled in blue and the small subunit in yellow. Scale bars, 150 nm in (a) and 150 nm in (b).

## DISCUSSIONS

In this work, we developed an efficient workflow VHUT-cryo-FIB combining vibratome, HPF, ultramicrotome pre-trimming and cryo-FIB milling for preparing high quality cryo-lamella of tissue specimen. In order to deal with various tissue samples, we introduced the vibratome section at the initial step. We tried to minimize the needs of equipment modification by only modifying the cryo-holder tip or the HPF carrier. Through this workflow, we have successfully fabricated cryo-lamella of skeleton muscle tissue from mice and the leaf tissue from spinach. The fabricated cryo-lamella can be further used for cryo-TEM imaging and high quality cryo-ET data collection. The reconstructed tomograms showed abundant structural features. Based on these tomograms, we further obtained *in situ* 3D structure of spinach ribosome that has never been reported before. All these results reveal a possibility of further use of our workflow for other *in situ* structural studies. However, there are still lots of improvements to make the workflow more productive, such as increasing the throughput of the procedure, adding more checkpoints during sample preparation, reducing the sample transfer operations to avoid ice contamination and combining correlative electron and light microscopy (CLEM) for locating target positions.

## MATERIALS AND METHODS

### Tissue sample acquirement and vibratome slicing

Muscle used in this study were obtained from mice aged 6-8 weeks, purchased from Institute of Experimental Animals, Chinese Academy of Medical Sciences. Mice were sacrificed by cervical dislocation and muscle was removed and cut into small (∼ 5 mm^3^) bulks with a razor blade. Then a bulk of muscle was washed 2 to 3 times in cold PBS and fixed on the specimen holder of VT1200S vibrating blade microtome (Leica Biosystems, US). The buffer tray of microtome was ice bathed and filled with cold PBS while slicing. The thickness of each slice was set to 50-100 µm depending on the cutting performance. The generated muscle slice was cut into appropriate size to fill into the HPF carrier.

For the spinach leaf specimen, we simply used a puncher to cut a circular slice at ∼ 2 mm diameter to fill into the HPF carrier.

### High pressure freezing

High pressure freezing was performed by HPF COMPACT 01 (Wohlwend Engineering, Switzerland). For the tissue sample in strategy A, the normal HPF carrier (3mm copper gold-plated from Leica microsystem with the product number #16770152) used is commercially available. For the tissue sample in strategy B, the specially designed carrier (**Supplementary Fig. 1**) was first assembled with the auxiliary ring before using. Tissue sample slice was put in the recess of the carrier and the cryoprotectant (1-hexadecene) was added to fill the surrounding area of the tissue slice. Then a sapphire disk was placed on the top of the carrier before the whole sandwich assembly was fast frozen by HPF. The frozen sample was removed from the clamping apparatus and stored in liquid nitrogen.

### Pre-trimming using cryo-ultramicrotome

After HPF the carrier was cryo-trimmed by Leica EM FC7 cryo-ultramicrotome (Leica microsystem, Austria) via a diamond knife (Diatome Trim 90, Switzerland) under -160 °C. The copper carrier was trimmed about one half by the diamond knife, leaving an embossment of 100 (width) × 20 (thickness) × 30 (depth) μm^3^ flat tissue specimen on the carrier (**Supplementary Movie 1**). Then the carrier was loaded onto a customized transfer shuttle ^17^. For Strategy B, the trimmed carrier was mounted onto FEI AutoGrid before loading.

### Cryo-FIB milling

Cryo-FIB milling was performed using a FIB/SEM dual beam microscope (Helios NanoLab 600i, Thermo Fisher Scientific, US) equipped with a Quorum PP3000T cryo-stage (Quorum Technologies, East Sussex, UK). The shutter with the carrier loaded was transferred into the FIB/SEM chamber by using Quorum PP3000T cryotransfer system (Quorum Technologies, East Sussex, UK) under -180 °C. To improve sample conductivity and reduce curtaining artifacts, the samples were deposited with organometallic platinum using the *in situ* gas injection system (GIS) operated at 5 seconds gas injection time before milling. During the milling process, the milling angle is nearly in parallel with the carrier. For the carrier loaded onto FEI AutoGrid in Strategy B, the cryo-stage was tilted a small angle of ∼ 10° to avoid beam blocked by the edge of AutoGrid. The milling was performed parallel at both sides of the sample platform to produce the cryo-lamella. Rough milling is performed with the accelerating voltage of the ion beam at 30 kV, and the current at 0.43 ∼ 0.79 nA. To facilitate tomography data collection, the notches surrounding the cryo-lamella were removed to yield a trapezoid shape (**Supplementary Movie 1**). After rough milling, one end of the lamella was jagged from the main platform. When the thickness of lamella reaches ∼1 μm, the ion beam current was reduced to 80 ∼ 230 pA to perform a fine milling. The final cryo-lamella with the thickness of 200 ∼ 300 nm was obtained.

### Cryo-TEM imaging and tilt series data collection

In Strategy A, a 200kV field emission cryoTEM microscope FEI Talos F200C (ThermoFisher Scientific, US) equipped with an FEI 4K × 4K Ceta camera (ThermoFisher Scientific, US) was used. The carrier containing the cryo-lamella was loaded onto a side entry cryo holder Gatan 910 (Gatan, US) with our customized holder tip (**Figs. 2e and 2g**). Then the carrier was inserted into the columne of the microscope for cryo-TEM imaging. The microscopy was operating at 200 kV. Cryo-TEM imaging was performed in a low dose mode with a defocus range of -8 ∼ -10 μm. The magnification was set up to yield a calibrated pixel size of 0.58 nm in the final recorded image.

In Strategy B, a 300kV field emission cryoTEM microscope FEI Titan Krios (ThermoFisher Scientific, US) equipped with a Gatan K2 Summit camera (Gatan, US) was used. The carrier containing the cryo-lamella had been preloaded onto FEI AutoGrid (**Fig. 2p**), which was loaded into the microscope by FEI Autoloader. The microscopy was operating at 300 kV. Cryo-TEM imaging was performed in a low dose mode with a defocus range of -4 ∼ -6 μm. The magnification was set up to yield a calibrated pixel size of 0.265 nm in the final recorded image.

SerialEM^18^ was used to collect tilt series data with the tilt angle from 40° to - 40° and the angle increment of 2°. The total dose used in a single tilt series was ∼ 100 e^-^/Å^2^. During data collection, two trial and autofocus positions were set 2 μm away at each side of the recording area in order to avoid redundant radiation damage.

### Image processing and sub-volume averaging

Dose fraction images recorded by Gatan K2 Summit camera were first motion corrected and summed by Motioncor2^19^, and then stacks were saved with the sequence based on the angle of each image by EMAN2.1 ^20^. For the subsequent sub-tomogram analysis, CTF estimation of each tilt image was performed by CTFFIND4 ^21^.

All the tilt series of images were aligned, reconstructed and post-processed by IMOD software package version 4.9.0 ^22^. Since there were no fiducial markers on the cryo-lamella, we selected patch tracking for image alignment. For the final reconstruction by weight back projection using IMOD ^22^, the radial filter options were set up with the cut-off of 0.35 and the fall-off of 0.05. The reconstruction of rat muscle sample was performed by using ICON ^23,24^ to get a better contrast for visualization.

From the tomogram of the spinach leaf specimen, ribosome particles were picked manually. Then the selected particles were extracted, classified and averaged by RELION1.4 ^25^. CTF model of each particle was generated by using RELION script.

In the present study, three tomograms from two different cryo-lamellas of spinach leaf were used for sub-tomogram analysis. Tomogram segmentation was performed using EMAN2.3 ^26^ and rendered with UCSF chimera (http://www.rbvi.ucsf.edu/chimera).

## DATA AVAILABILITY

The cryoET tomograms of rat skeleton muscle and spinach leaf have been deposited into Electron Microscope Data Bank (EMDB) with the accession codes EMD-XXXX and EMD-XXXX, respectively. The corresponding data sets of raw cryoET tilt series have been deposited into EMPIAR with the accession codes EMPIAR-XXXX and EMPIAR-XXXX, respectively. The sub-tomogram averaged map of spinach ribosome was deposited into EMDB with the accession code EMD-XXXX.

## AUTHOR CONTRIBTUIONS

F. S. initiated and supervised the project. J.Z., D.Z. and L.S. performed all the experiments. J.Z. and G.J. made initial design of carriers. X.H. wrote SerialEM scripts for cryo-electron tomography data collection. D.Z and N.T. performed image processing including reconstruction and sub-tomogram averaging. J.Z., D.Z. and F.S. analyzed the data and wrote the paper.

## ACKNOWLEDGMENTS

We are grateful to Xinyi Wu and Sai Yang from Laboratory Animal Research Center, Institute of Biophysics, Chinese Academy of Sciences, for providing available male mice. We would like to thank Prof. Wanzhong He and Dr. Zhaodi Jiang from Beijing National Institute of Biological Science for their help on high pressure freezing. We would also like to thank Ping Shan and Ruigang Su (F.S. lab) for their assistances of lab management.

This work was supported by grants from National Natural Science Foundation of China (31830020) and the Ministry of Science and Technology of China (2017YFA0504702). All the sample preparation and cryoEM works were performed at Center for Biological Imaging (CBI, http://cbi.ibp.ac.cn), Institute of Biophysics, Chinese Academy of Sciences.

## COMPLIANCE WITH ETHICAL STANDARDS

All authors declare that they have no conflict of interest. All institutional and national guidelines for the care and use of laboratory animals were followed.

## SUPPLEMENTARY FIGURES AND MOVIES

**Figure S1.**
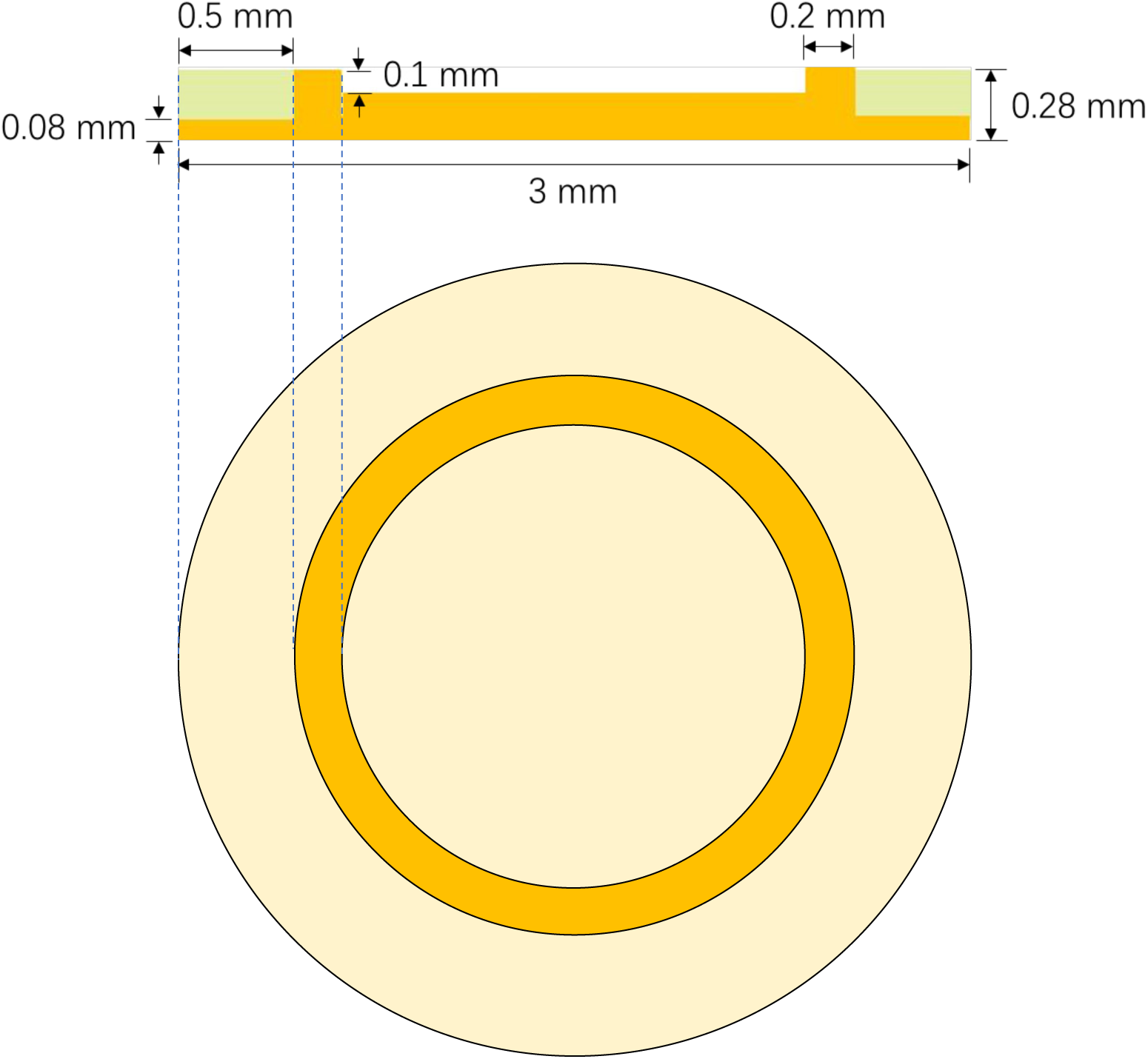
Detailed design of the modified carrier used in strategy B. The auxiliary ring is colored in light green.

**Figure S2.**
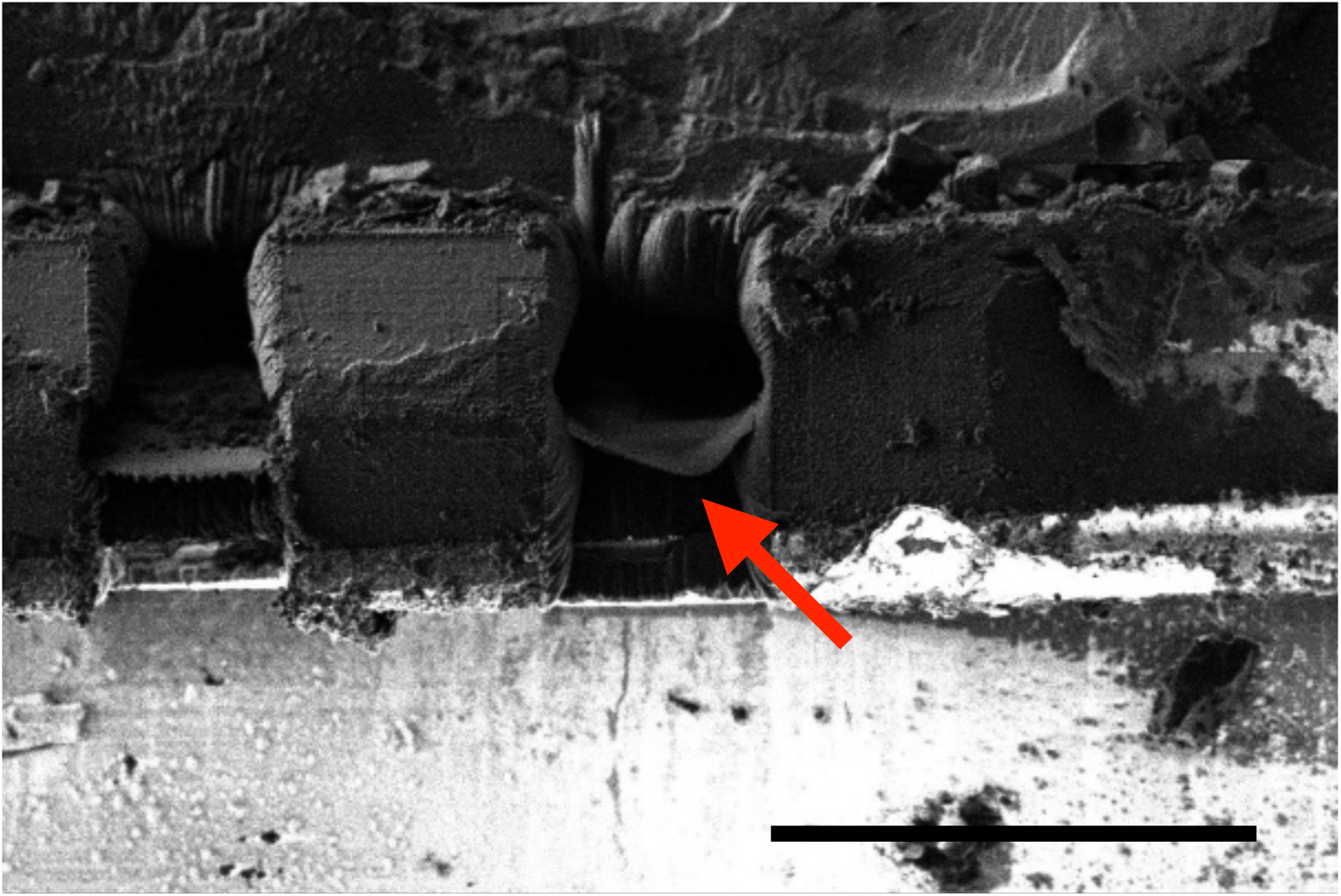
Observation of bending of cryo-lamella during cryoFIB milling. Secondary electron micrograph in FIB mode shows a bend milled cryo-lamella that is indicated by the red arrow. Scale bar, 50 μm

**Figure S3.**
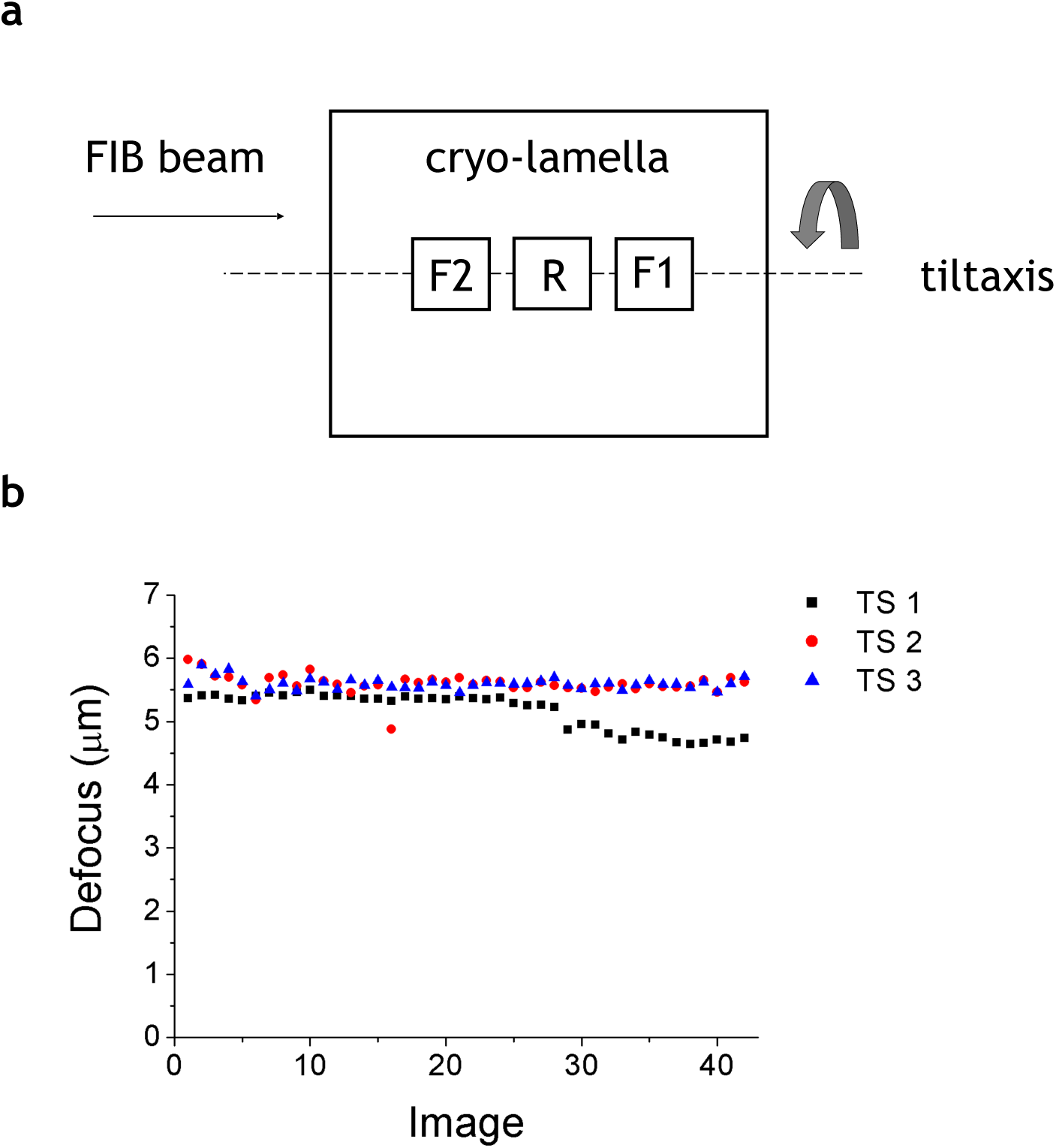
Cryo-ET data collection of cryo-lamella. (a) Scheme of multi-focus strategy used for tilt series data collection. R, record (exposure); F, focus.Defocus distribution of three tilt series collected using multi-focus strategy.

**Figure S4.**
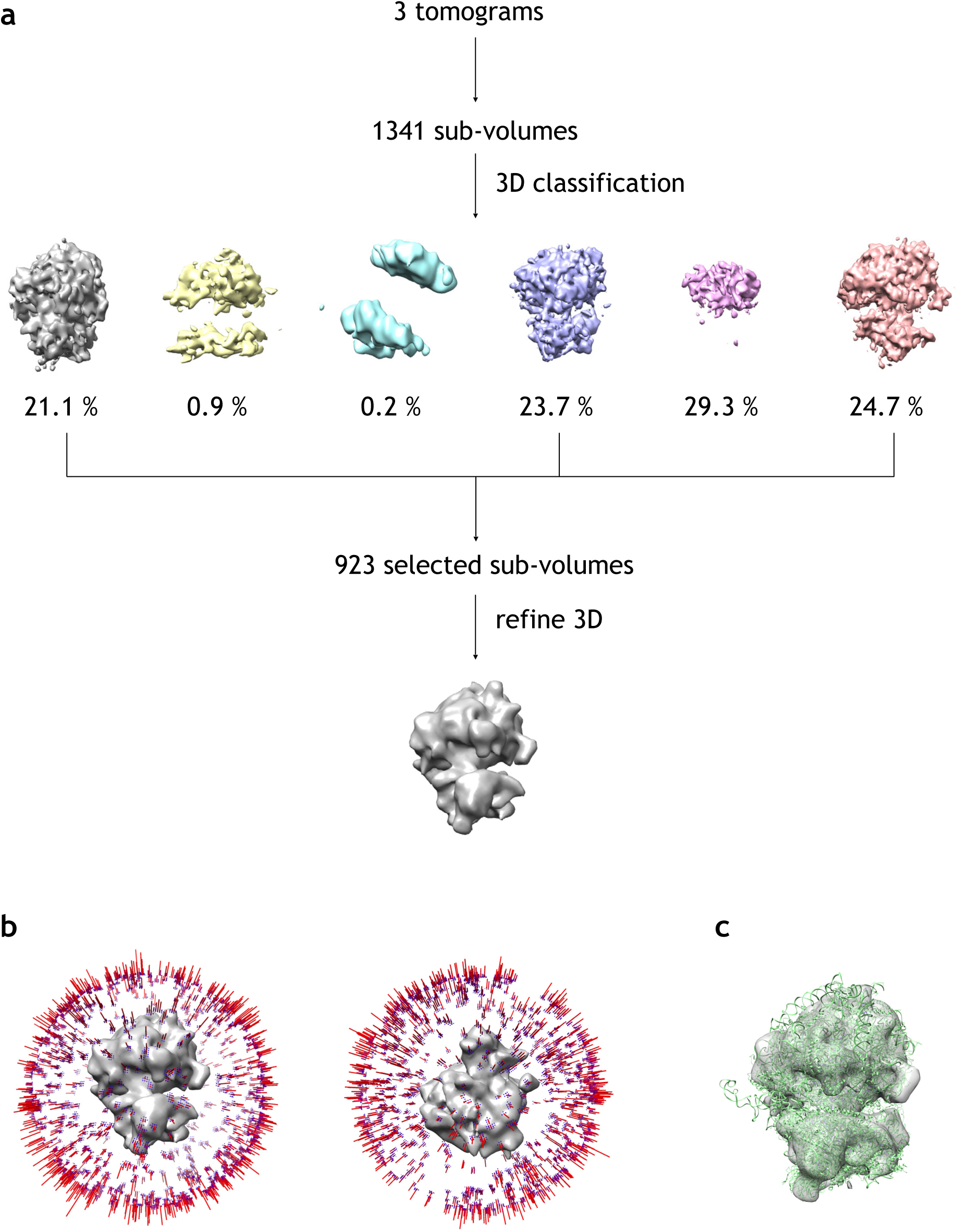
Procedure of sub-tomogram averaging. (a) Workflow of image processing. (b) Final averaged map and the angular distribution of the sub-tomogram particles. (c) Fitting human 80S ribosome structure (PDB ID 6EK0, colored in green) into the sub-tomogram averaged map.

**Movie S1. A complete VHUT-cryo-FIB workflow in Strategy B.**

**Movie S2. Aligned cryo-ET tilt series of cryo-lamella of rat skeleton muscle specimen.**

**Movie S3. Slicing view of tomogram of cryo-lamella of rat skeleton muscle specimen.**

**Movie S4. Aligned cryo-ET tilt series of cryo-lamella of spinach leaf specimen.**

**Movie S5. Slicing view of tomogram of cryo-lamella of spinach leaf specimen.**

